# Optimal Mass Transport for Robust Texture Analysis

**DOI:** 10.1101/855221

**Authors:** Zehor Belkhatir, Aditi Iyer, James C. Mathews, Maryam Pouryahya, Saad Nadeem, Joseph O. Deasy, Aditya P. Apte, Allen R. Tannenbaum

## Abstract

The emerging field of radiomics, which consists of transforming standard-of-care images to quantifiable scalar statistics, endeavors to reveal the information hidden in these macroscopic images. This field of research has found different applications ranging from phenotyping and tumor classification to outcome prediction and treatment planning. Texture analysis, which often consists of reducing spatial texture matrices to summary scalar features, has been shown to be important in many of the latter applications. However, as pointed out in many studies, some of the derived texture statistics are strongly correlated and tend to contribute redundant information; and are also sensitive to the parameters used in their computation, e.g., the number of gray intensity levels. In the present study, we propose first to consider texture matrices, with an emphasis on gray-level co-occurrence matrix (GLCM), as a non-parametric multivariate objects. The proposed modeling approach avoids evaluating redundant and strongly correlated features and also prevents the feature processing steps. Then, via the Wasserstein distance from optimal mass transport theory, we propose to compare these spatial objects to identify computerized tomography slices with dental artifacts in head and neck cancer. We demonstrate the robustness of the proposed classification approach with respect to the GLCM extraction parameters and the size of the training set. Comparisons with the random forest classifier, which is constructed on scalar texture features, demonstrates the efficiency and robustness of the proposed algorithm.

## I. INTRODUCTION

The importance of medical imaging has expanded to all major area of health care, including oncology. Motivated by the goal of unveiling the information hidden in standard-of-care images, the research field of radiomics has emerged [1]. Radiomics, which consists of the high-throughput extraction of quantitative features from medical images using automated quantification algorithms, is becoming a powerful tool in different applications, e.g., outcome prediction, tumor classification, treatment planning, and personalized therapy [2], [3], [4], [5] to cite few examples. Texture analysis, which refers to the quantification of spatial variations in gray levels within an image or region of interest (ROI), is widely used and has been successful in different studies, e.g., [6], [7], [8].

Usually in the radiomics research field, characterizing texture consists first of computing multivariate texture matrices such as gray-level co-occurrence matrix (GLCM), run-length matrix (RLM), size zone matrix (SZM), and neighborhood gray-tone difference matrix (NGTDM), and then reducing these multivariate objects to scalar summary statistics. This reduction step may lead to a loss of the spatial information that is inherent in texture matrices. In fact, Vickers and Modestino [9] noted in their work that using scalar features of GLCM for a classification task is suboptimal and that better results may be obtained by using the spatial GLCM directly in their proposed maximum likelihood classifier. Moreover, it has been pointed out in several studies that some of the resulting summary statistics (i) are highly correlated which may lead to overfitting [10], [11], [12], and (ii) also depend on gray-level discretization [13]. In [14], the authors propose to model GLCM as a multivariate object using a latent Gaussian Markov random field model in order to avoid the loss of spatial information. The method has shown promising results in classifying benign and malignant adrenal lesions. Moreover, considering the GLCM as a density function resulted in texture features that are less sensitive to the gray-level quantization, as shown in [15], [16].

Motivated by these recent works, we propose in this study: (i) to consider texture matrices in general, and GLCM in particular, as non-parametric multivariate objects; and (ii) use Wasserstein distance, from Optimal Mass Transport (OMT) theory, as a metric to compare them for the purpose of classifying computed tomographic (CT) slices with dental artifacts in head and neck cancer (H&NC). The OMT problem seeks the most efficient way to transform one distribution of mass to another given a cost function [17]. It origins go back in 1781, when Gaspar Monge formulated the problem of finding the minimal transportation cost to redistribute earth for building fortifications [18]. Leonid Kantorovich in 1942, relaxed Monge’s formulation to find an optimal coupling of distributions using linear programming [19]. Since then, OMT has played a crucial role in many fields of science and engineering; see [20], [21]and references therein. The increased interest in the use of OMT-based metrics, known as Wasserstein distance or Earth-Mover’s-Distance (EMD) in the engineering field, is mainly due to their natural ability to capture spatial information when comparing signals, images, or other types of data. This allows to provide various data distributions with different geometric interpretations, which we are seeking to capture from spatial texture matrices in the present work.

## II. MATERIALS AND METHODS

Dental artifacts can dramatically attenuate the quality of CT images and make their analysis challenging. It is essential, therefore, to develop an automated algorithm to detect CT slices with dental artifacts in H&NC. This preliminary preprocessing phase will allow taking measures to reduce the effect of the noisy slices when using the images for other modeling purposes. In this section, we present an automatic classification algorithm, which is based on the idea of considering GLCM as a multivariate object, not a summary of scalar statistics, and employs the Wasserstein distance to compare pairs of GLCMs.

### A. Data characteristics

The proposed method was applied to two datasets: (i) an internal dataset of 44 H&N CT scans from our institution, resulting in 1165 axial slices with voxel sizes ranging from 0.0914 − 0.1367 cm (median: 0.0977 cm) and slice thickness 0.1 − 0.3 cm (median: 0.25 cm), and (ii) an external dataset of 24 H&N CT scans from the open-source archive TCIA [22], resulting in 679 slices with voxel sizes ranging from 0.0352 − 0.0977 cm (median: 0.0488 cm) and slice thickness 0.0450 − 0.5009 cm (median: 0.25 cm). Both datasets were resampled to 0.0977 cm× 0.0977 cm× 0.25 cm prior to analysis, resulting in 1235 and 620 slices from internal and external datasets, respectively. Each CT slice was labeled as noisy or clean based on the presence of dental artifacts by a medical imaging expert, resulting in 334 (27.04%) noisy and 901 (72.96%) clean slices from MSK cohort, and 53 (8.55%) noisy and 567 (91.45%) clean slices from TCIA cohort.

### B. Radiomic texture features

CT images were thresholded (at the 5th percentile) to exclude regions of air, followed by morphological processing to extract the patient’s outline. GLCMs were extracted within the resulting mask using the radiomics extension of the Computational Environment for Radiological Research (CERR) [23]. GLCMs were computed by combining contributions from all 2-D neighbors, using 4 different gray levels from 16 − 128 and 5 different sampling rates between ±15% from the original resolution. The resulting parameters are listed in Table I. Additionally, 25 scalar features, which are listed in Table II, were extracted from the GLCMs to be used in a machine learning classifier for comparison purpose.

**TABLE I:** GLCM extraction parameters.

|  |  |
| --- | --- |
| Voxel resolution | 0.0830 cm × 0.0830 cm, 0.0903 cm × 0.0903 cm, 0.0977 cm × 0.0977 cm, 0.1050 cm × 0.1050 cm, 0.1123 cm × 0.1123 cm |
| No. gray levels | 16, 32, 64, 128 |

**TABLE II:** GLCM-based scalar features.

|  |  |  |
| --- | --- | --- |
| 1 : Energy | 10 : Inverse difference normalized | 18 : Cluster tendency |
| 2 : Joint entropy | 11 : Inverse variance | 19 : Cluster shade |
| 3 : Joint max | 12 : Difference entropy | 20 : Cluster prominence |
| 4 : Joint average | 13 : Difference variance | 21 : Haralick correlation |
| 5 : Joint variance | 14 : Sum average | 22 : Auto-correlation |
| 6 : Contrast | 15 : Sum entropy | 23 : First measure of information correlation |
| 7 : Inverse difference moment | 16 : Sum variance | 24 : Second measure of information correlation |
| 8 : Inverse difference moment normalized | 17 : Correlation | 25 : Dissimilarity |
| 9 : Inverse difference |  |  |

### C. Image classification algorithm using spatial GLCMs and Wasserstein distance

We use the theory of OMT [17] to define distances between spatial GLCMs, which we consider as 2-D probability distributions.

Let *ρ*_0_, *ρ*_1_ be two *d*-dimensional probability distributions defined on Ω ⊂ ℝ^*d*^. The ***Wasserstein-1 distance*** is defined as follows:

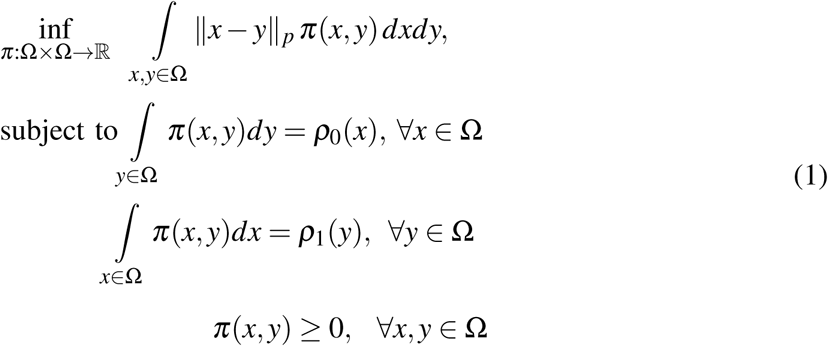

where ‖.‖_*p*_, 1 ≤ *p* ≤ ∞, is the ground metric of the Wasserstein distance. The variable *π* is the set of joint distributions *π* : Ω × Ω → ℝ whose marginal distributions are *ρ*_0_, *ρ*_1_.

An equivalent alternative formulation of the Wasserstein-1 distance, which is simpler and computationally more efficient, is defined by the following optimization problem:

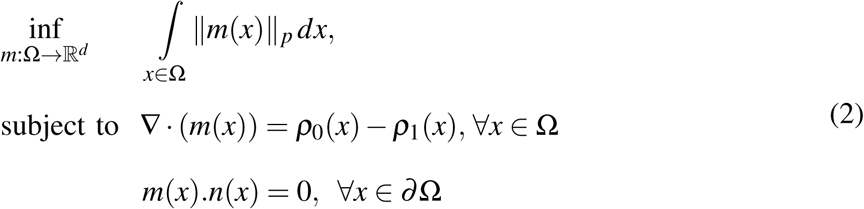

where “∇·” denotes the divergence operator, *n*(*x*) is the normal to the boundary *∂*Ω, and *m* is a *d*-dimensional field satisfying the zero flux boundary condition [24]. A fast numerical scheme that relies on multilevel primal-dual optimization algorithms was proposed in [25] to solve (2). This latter numerical scheme is adopted in the present study.

#### Proposed classification algorithm

We start by normalizing the computed GLCMs to be 2-D probability distributions. This step, essential to apply the balanced OMT theory introduced earlier, doesn’t affect the textural pattern. However, interestingly, it was found in recent studies that this step offers some advantages. It adjusts for heterogeneity in lesion size [14], and produce texture features that are invariant to the quantization gray-levels [15], [16]. We then partition the available data to training and test sets, where the training dataset consists of a subset of MSK slices, and the test dataset is comprised of the whole TCIA slices. The proposed supervised classifier mainly relies on comparing 2-D normalized GLCMs of the test CT slices with the ones of the training set using Wasserstein-1 distance. We point out that the comparison between the GLCMs extracted from MSK and TCIA cohorts is performed using the same values of the extraction parameters (voxel resolution and number of gray levels). The main steps of the proposed classification algorithm, which are depicted in Figure 1, are given as follows

**Fig. 1:**
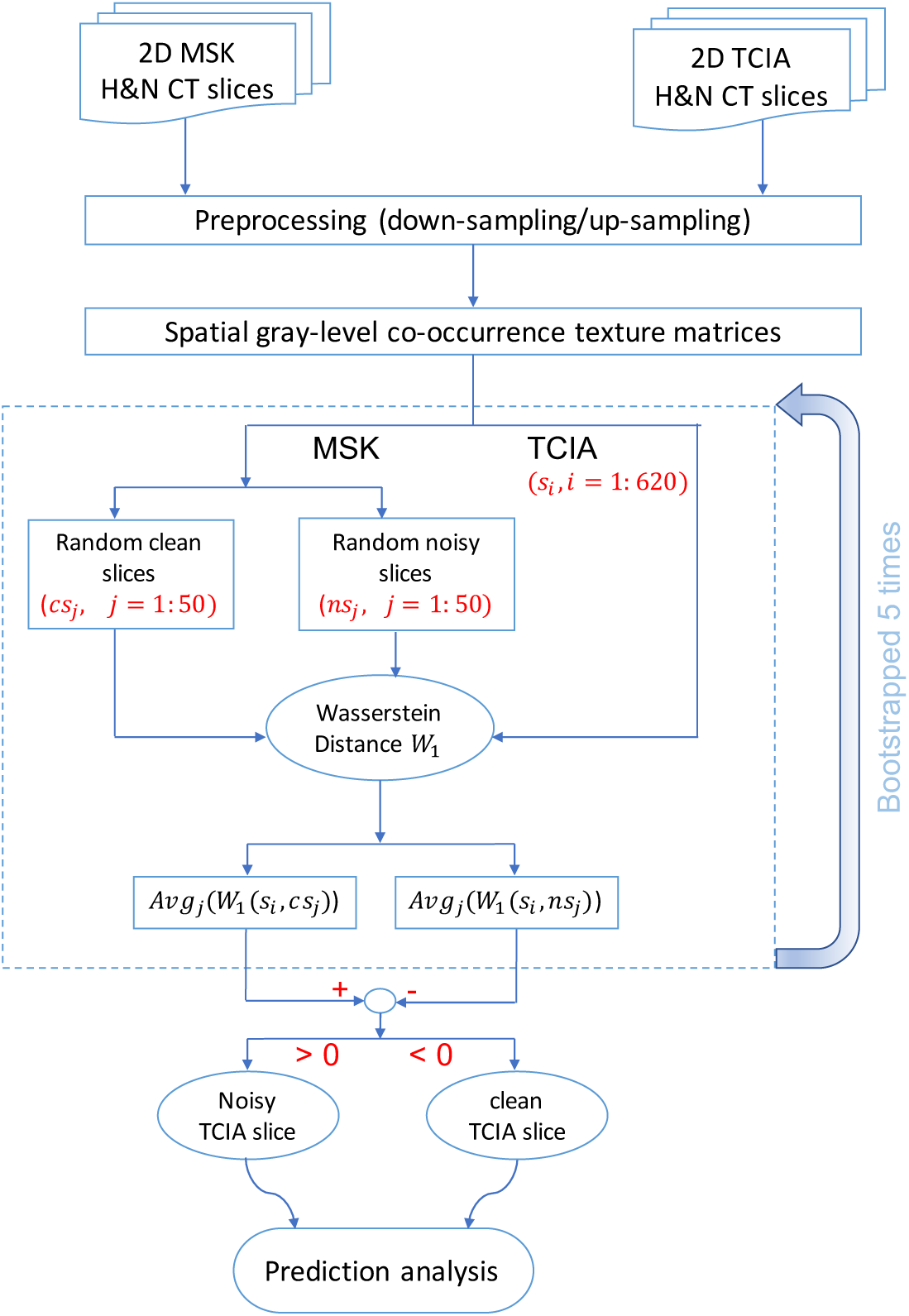
Flowchart of the proposed classification algorithm for CT slices with dental artifacts in H&NC.

i. We randomly select a subset, *N*_1_ = 50, of noisy and clean slices, denoted *ns*_*j*_ and *cs* _*j*_, for *j* = 1 : *N*_1_, respectively, from the training cohort.
ii. Then, for a given TCIA slice denoted *s*_*i*_, *i* = 1 : *N*_2_, with *N*_2_ = 620, we compute the Wasserstein distance between its corresponding GLCM and the ones of all noisy and clean MSK slices. This step is repeated 5 times, resulting two matrices 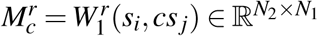 and 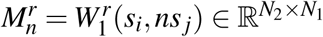, for *r* = 1 : 5.
iii. Based on the averaged comparison with respect to the training dataset between 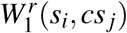 and 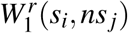, the class for slice *s*_*i*_ is concluded.
iv. Finally, Youden’s J index = (sensitivity + specificity – 1), which is a way of summarizing sensitivity and specificity metrics into a single numeric value, was used to assess the accuracy of the classification results.

## III. RESULTS

We evaluated the performance of the proposed algorithm for different parameter combinations involved in the computation of GLCM. We also compared the results of the proposed classifier with the random forest classifier, which uses the scalar features shown in Table II. Robustness of both algorithms with respect to the GLCM computation settings and size of the training set was carried out as well.

Figure 2 depicts Youden’s J index that combines the performances of classifying the noisy and clean slices correctly, i.e., sensitivity and specificity, in a single number. The best performance of 77.76% (sensitivity: 87.2%, specificity: 90.57%) is achieved for a gray-level discretization level of 16 and voxel resolution of 0.83 mm × 0.83 mm. It is worth noting, however, that the performance for other combination pairs of GLCM extraction parameters is also good with small variability between the relatively bad and best performances (mean = 76.66 %, std = %). Regarding the random forest classifier, the best performance of 76.24% (sensitivity: 89.07%, specificity: 87.17%) is achieved with gray-level quantization of 32 and voxel resolution of 1.05 mm × 1.05 mm as illustrated in Figure 2. We emphasize, however, that the performance of the random forest technique is more sensitive to the GLCM computation settings (mean = 69.94 %, std = 3.63 %).

**Fig. 2:**
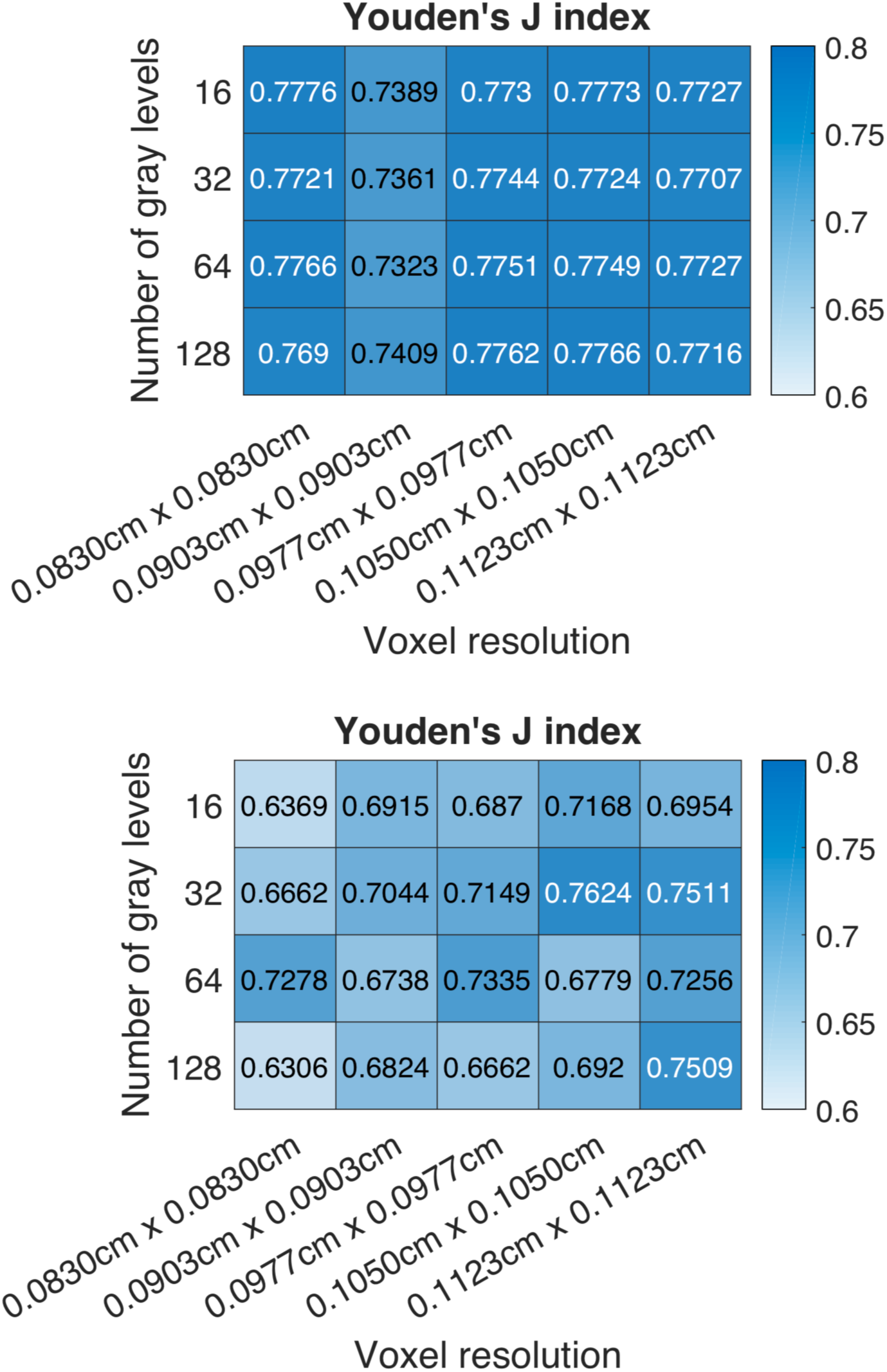
Classification accuracy results using 100 traning data (50 noisy and 50 clean). Upper panel: Proposed OMT/spatial GLCM classifier. Lower: Random forest classifier.

We also tested the sensitivity of the performance of the classifiers to different sizes of training data. The results of the proposed classifier and the random forest are given in Tables III and IV, respectively. The OMT/spatial GLCM based method performs better than the random forest approach for a small number of training data (10 slices). Even for this small number of training data, the proposed algorithm was not very sensitive to the GLCM computation settings, while the random forest method fails to achieve good performance for all the considered extraction parameter pairs, as depicted in Figure 3. Increasing the size of the training set leads to a significant increase in the accuracy of the random forest approach, while the proposed method achieves comparable good classification results (see Tables III and IV).

**TABLE III:**
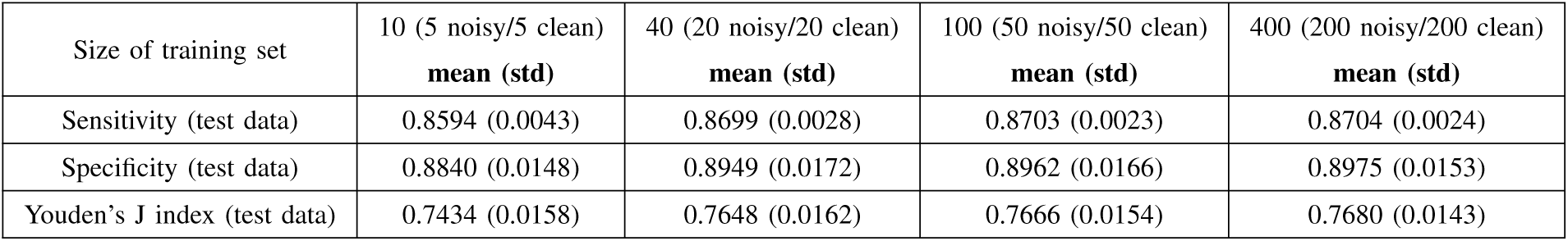
Robustness of the classification results to the size of training dataset using the proposed OMT/spatial GLCM based classifier. The **mean** and **std** represent the average and standard deviation of the classification accuracy for the different combination pairs of GLCM extraction parameters.

| Size of training set | 10 (5 noisy/5 clean)<br>**mean (std)** | 40 (20 noisy/20 clean)<br>**mean (std)** | 100 (50 noisy/50 clean)<br>**mean (std)** | 400 (200 noisy/200 clean)<br>**mean (std)** |
| --- | --- | --- | --- | --- |
| Sensitivity (test data) | 0.8594 (0.0043) | 0.8699 (0.0028) | 0.8703 (0.0023) | 0.8704 (0.0024) |
| Specificity (test data) | 0.8840 (0.0148) | 0.8949 (0.0172) | 0.8962 (0.0166) | 0.8975 (0.0153) |
| Youden's J index (test data) | 0.7434 (0.0158) | 0.7648 (0.0162) | 0.7666 (0.0154) | 0.7680 (0.0143) |

**TABLE IV:**
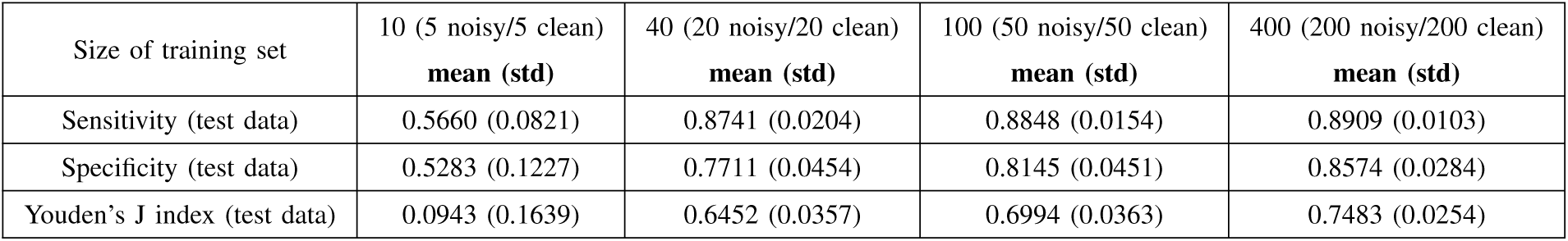
Robustness of the classification results to the size of training dataset using random forest and GLCM scalar features. The **mean** and **std** represent the average and standard deviation of the classification accuracy for the different combination pairs of GLCM extraction parameters.

**Fig. 3:**
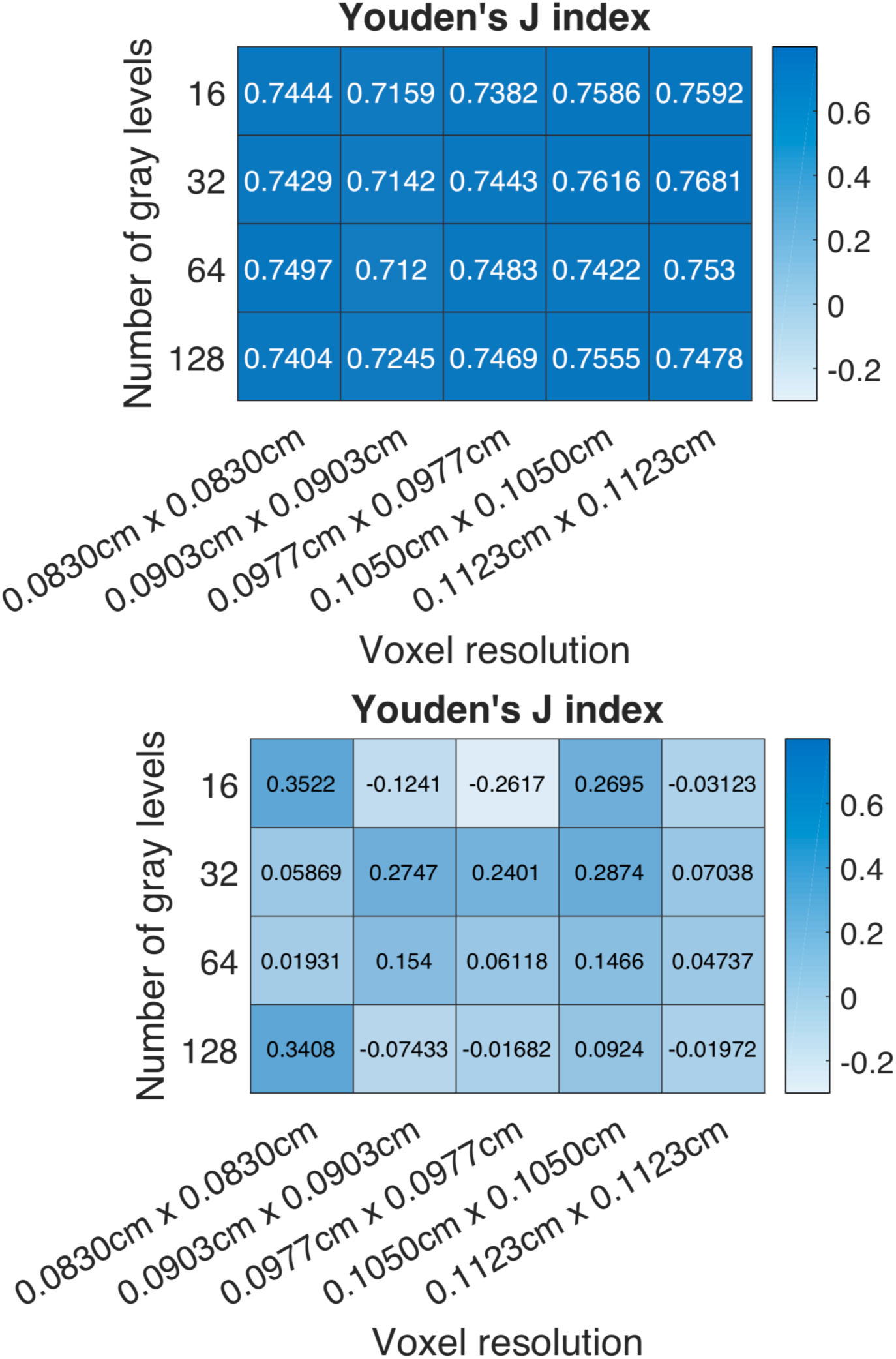
Classification accuracy results using 10 traning data (5 noisy and 5 clean). Upper panel: Proposed OMT/spatial GLCM classifier. Lower: Random forest classifier.

## IV. DISCUSSION

In this study, we investigated two ideas related to texture analysis for medical images. The first idea is based on directly utilizing texture matrices, not their scalar summary statistics, as multivariate radiomic features. The second idea consists of using a robust metric, the so-called Wasserstein distance, on these high-dimensional objects to classify CT images with dental artifacts in H&NC. Combining these two ideas has resulted in a classification algorithm that is less sensitive to the GLCMs computation parameters. It also does not need a large number of training data to achieve high classification performance.

The potential of using the GLCM as a multivariate object for an oncological application recently introduced in [14]. In this work, the authors used a Gaussian Markov random field model to characterize GLCMs. Their modeling approach was employed to distinguish between benign and malignant adrenal lesions based on CT scans, and the obtained results were promising compared to different machine learning methods that were based on scalar feature quantities. The concept though of using multidimensional parametric models such as Markov random field to capture texture is old [26], [27], but never used in oncology to the best of our knowledge.

Even though there is no precise universal definition for texture [11], it still remains a key element of visual perception to analyze images in different fields, particularly the cancer research field. Therefore, more efforts need to be employed to go beyond the summary texture features by considering the multivariate structure of texture matrices. Doing so, hopefully, will avoid information loss of the functional patterns that describe the perfusion, density, or morphology of tumor microenvironment.

This work is the first proof-of-concept study that offers some new insights in dealing with texture to reveal the intrinsic heterogeneity observed in medical images. Further research investigations need to be conducted in the future to show the applicability and potential use of multivariate texture matrices (GLCM, RLM, SZM, NGTDM) and OMT theory for other purposes besides medical image classification, e.g., in outcome prediction and treatment planning. A potential future direction is to combine the geometrical texture features, which consists of the Wasserstein distance between texture lattices, with other types of radiomic features, e.g., shape, to build appropriate machine learning prediction models.

## V. CONCLUSION

We developed a classification algorithm for 2D medical images based on the Wasserstein distance and the multivariate GLCMs. Overall, the proposed classifier approach outperformed the random forest method, which does not leverage the spatial information in GLCMs, in terms of robustness to the GLCM extraction parameters as well as the size of the training set.

## ACKNOWLEDGMENTS

This study was supported by AFOSR grant (FA9550-17-1-0435), a grant from National Institutes of Health (R01-AG048769), MSK Cancer Center Support Grant/Core Grant (P30 CA008748), and a grant from Breast Cancer Research Foundation (BCRF-17-193).

